# Somatic Loss of the Y Chromosome and Alzheimer’s Disease Risk

**DOI:** 10.1101/2022.11.14.516433

**Authors:** Ellen Palmer, Penelope Benchek, Nicholas Wheeler, Sandra Smeiszek, Adam C. Naj, Jonathan L. Haines, Margaret A. Pericak-Vance, Lars A. Forsberg, Holly N. Cukier, Yeunjoo Song, William S. Bush

## Abstract

Mosaic loss of the Y chromosome (LOY) is a somatic, age-related event that has been previously associated with a variety of diseases of aging. A prior study of European cohorts demonstrated an association between LOY and Alzheimer’s Disease and more recent molecular studies have shown that LOY can also occur within microglia, suggesting a potential functional role in AD pathogenesis. In this study, we further validate the association between LOY and AD via prospective analyses of 1,447 males, and perform Mendelian Randomization analysis on 10,013 males across 26 US cohorts. Significant results from these analyses provide further evidence for a role of LOY in the development of Alzheimer’s Disease.

## Introduction

Alzheimer’s disease (AD [MIM: 104300]) is a complex, multifactorial disorder of aging that results in both structural and functional changes to the brain. Since the early 90’s, multiple germline genetic factors have been associated to AD risk [1]. Many of these genetic factors are associated with the abnormal production and accumulation of amyloid beta and tau proteins [1], which leads to plaques, neurofibrillary tangles, neuronal death, and loss of brain cortical volume [2]. Bolstered by the association of rare genetic coding variants in the *TREM2* gene [3–6], the clearance of amyloid plaques by microglia has emerged as an important disease mechanism [7]. With this and subsequent genetic associations, immune system function is now thought to play a critical role in AD development [8].

Due to its limited genetic content and lack of recombination outside of the pseudoautosomal regions, the Y chromosome has rarely been the focus of genomic studies. However, early in the study of cytogenetics, karyotypes of peripheral immune cells revealed occasional somatic loss of the Y chromosome (LOY) in males [9]. Starting in 2014, multiple reports characterized LOY via intensity analysis of genotyping microarrays across cohorts containing men of advanced age within multiple European-descent cohort studies, first reporting associations of LOY to cancer risk [10]. Subsequent studies demonstrated an increased risk of incident AD diagnosis above a threshold of peripheral loss of Y [11], though the underlying mechanism that results in LOY remains unknown.

Multiple genetic association studies have examined loss of Y as an outcome. Through a study of 85,541 men, Wright et al. revealed 19 genomic regions and 36 differentially methylated sites associated with LOY, and containing or regulating genes whose functions converge on mechanisms of cell proliferation and cell cycle regulation [12]. These variants also predict X chromosome loss in women. Thompson et al. performed similar analyses in nearly one million samples to identify 156 autosomal genetic variants and indels [13]. Taken together, these data suggest that LOY in the peripheral blood is a potential proxy trait for estimating rates of mitotic aneuploidy, cell cycle efficiency, and genomic instability due to aging [10, 11]. More recent molecular work also suggests that LOY itself may drive phenotypic changes [14].

In 2016, Dumanski et al. associated mosaic LOY in blood with Alzheimer’s disease in one case/control (adjusted odds ratio = 2.80) and two prospective cohort studies [11]. More recently, Vermeulen et al. evaluated aged human brains via single-cell and single nuclei RNA-seq studies and found that LOY occurs in approximately 7.2% of microglia compared to 0.3% of neurons and 1.1% of astrocytes in the samples [15]. The rapid turnover of microglia may influence their susceptibility to LOY as compared to terminally differentiated cells [16]. Intriguingly, examination of differentially expressed genes in the microglia with LOY identified 193 genes, including the AD risk genes *APOE, ABCA1*, and *RHOH*[17, 18]. In addition, the study reported a significant increase in LOY of microglia from male AD donors.

Here, we seek to further understand the relationship between somatic loss of Y and AD across multiple US cross-sectional studies as part of the Alzheimer’s Disease Genetics Consortium (ADGC). We also perform Mendelian randomization to provide further evidence of the antecedence and potentially causal role of loss of Y in AD pathogenesis. Finally, we identify genetic variants distinct to LOY AD cases, providing the first evidence that LOY may be a marker of a different disease etiology.

## Methods

### Cohorts

Loss of Y chromosome estimation was conducted across multiple cohorts in the Alzheimer’s Disease Genetics Consortium. These included the Adult Changes in Thought study (ACT), the National Institute on Aging Alzheimer Disease Centers (ADCs), the Chicago Health and Aging Project (CHAP), the NIA-LOAD study, the Religious Orders Study/Memory and Aging Project (ROSMAP), and the Washington Heights-Inwood Community Aging Project (WHICAP). These cohorts were selected based on the availability of age at blood sampling and the use of genotyping arrays with sufficient probe coverage of the Y chromosome. Details of individual AD study designs, ascertainment, and phenotype definitions are provided in supplemental materials.

### Loss of Y Chromosome Estimation

Three genotyping platforms were used across the ADGC cohorts with sufficient Y chromosome probes to allow copy number determination (Illumina 610, 1M duo, OmniExpress or the HumanOmni1-Quad). Raw intensity data was accessed for all participants of each study. Using vendor-provided annotation files, we remapped all Y chromosome probe sequences to GRCh build 38 using BLAT and excluded probes with non-Y target effects. We also restricted samples to males only as assessed by X chromosome zygosity. We defined a continuous estimate of LOY mosaicism by computing the median of the log R ratio (mLRRY), calculated as the median value of probe intensities from the male specific portion of the chromosome Y relative to probes from chromosome 1. To eliminate systematic bias in the mLRRY distributions, within each cohort we implemented the experimental noise method as described in Forsberg et al. [10] figure 2. An mLRRY value close to zero indicates a normal chromosome Y state and more negative mLRRY values indicate increasing degree of LOY mosaicism. We also defined a binary LOY phenotype in two ways using both a 99% confidence limit cut-off and an estimated experimental variation cut-off. This approach has been applied and experimentally validated previously [11].

### Statistical Analyses

Joint analyses over all cohorts relating LOY estimates to AD case/control status were performed using a non-parametric Kolmogorov-Smirnov test. Adjusted analyses were performed using analysis of covariance (ANCOVA). Genome-wide association study analyses for LOY and AD outcomes were performed using Hail (version 0.1-74bf1eb) running on Apache Spark version 2.1.0, using linear and logistic regression. Betas and standard errors were extracted for use in Mendelian randomization analyses. Meta-analysis was conducted using regression models within each cohort followed by fixed-effects meta-analysis both with and without covariant adjustments. Meta-analyses were performed using METAL software, and the *metafor* R package.

### Mendelian Randomization and Genetic Risk Score Analyses

Unadjusted LOY betas and standard errors (SEs) were calculated using linear regression in the subset of biologically male samples where mLRRY was estimated. These values were compared to published betas by Wright et al. (Supplemental data) to ensure the published instrument was directionally concordant with the effect seen in our data, and thus a valid instrument. Due to differences in genotyping arrays and imputation quality between studies, we had 17 of the 19 SNPs published by Wright et al. for use as instruments.

We tested these same SNPs for AD associations using logistic regression models, adjusted for *APOE4* dose, testing the association between each SNP and AD status within each cohort. We then used the R package *metafor* to perform a fixed-effects meta-analysis. The betas and SEs for each SNP from the meta-analysis were retained for use in Mendelian Randomization (MR) models. These models were completed for a) the subset with measured LOY, b) all males, c) all females, and d) all ADGC samples to check for both sex-specific effects as well as an expected bias towards the null when using non-overlapping samples.

We ran MR models for the Wright instrument. The MR method requires as input the betas and SEs for associations to the instrument and outcome phenotypes, in this case LOY and AD respectively. All models were computed using the LOY betas and SEs calculated in our LOY-measured subset as the instrument. These were paired with each of the four AD betas and SEs for bias and sex-specific assessment. Models were run using the inverse-variance weighted meta-analysis (IVW) method. Finally, we performed a leave-one-out permutation sensitivity analysis to ensure the observed effects were not driven by a single variant. All analyses were completed in R using the *MendelianRandomization* package.

## Results

### Validation of Mosaic Loss of Y as an AD Risk Factor

Our QC process removed between 29 and 37% of poorly mapped chromosome Y probes among the different genotyping arrays which reduced noise and Y chromosome intensity outliers. 4700 males passed our QC process and had an available age at blood draw, with 633 (13.4%) showing LOY (using the 99% CI threshold approach). Of these, 3117 had a clear AD diagnosis as either case or control with 438 (14.1%) showing LOY. Age at blood draw was stratified into four bins consistent with prior literature (<=65, 66-75, 76-85, >=86), and Wilcoxan rank sum tests were performed to examine the distribution of LOY by age at blood sampling. This trend was highly significant over all samples and between all age groups, consistent with multiple prior reports (**Figure 1**).

**Figure 1.**
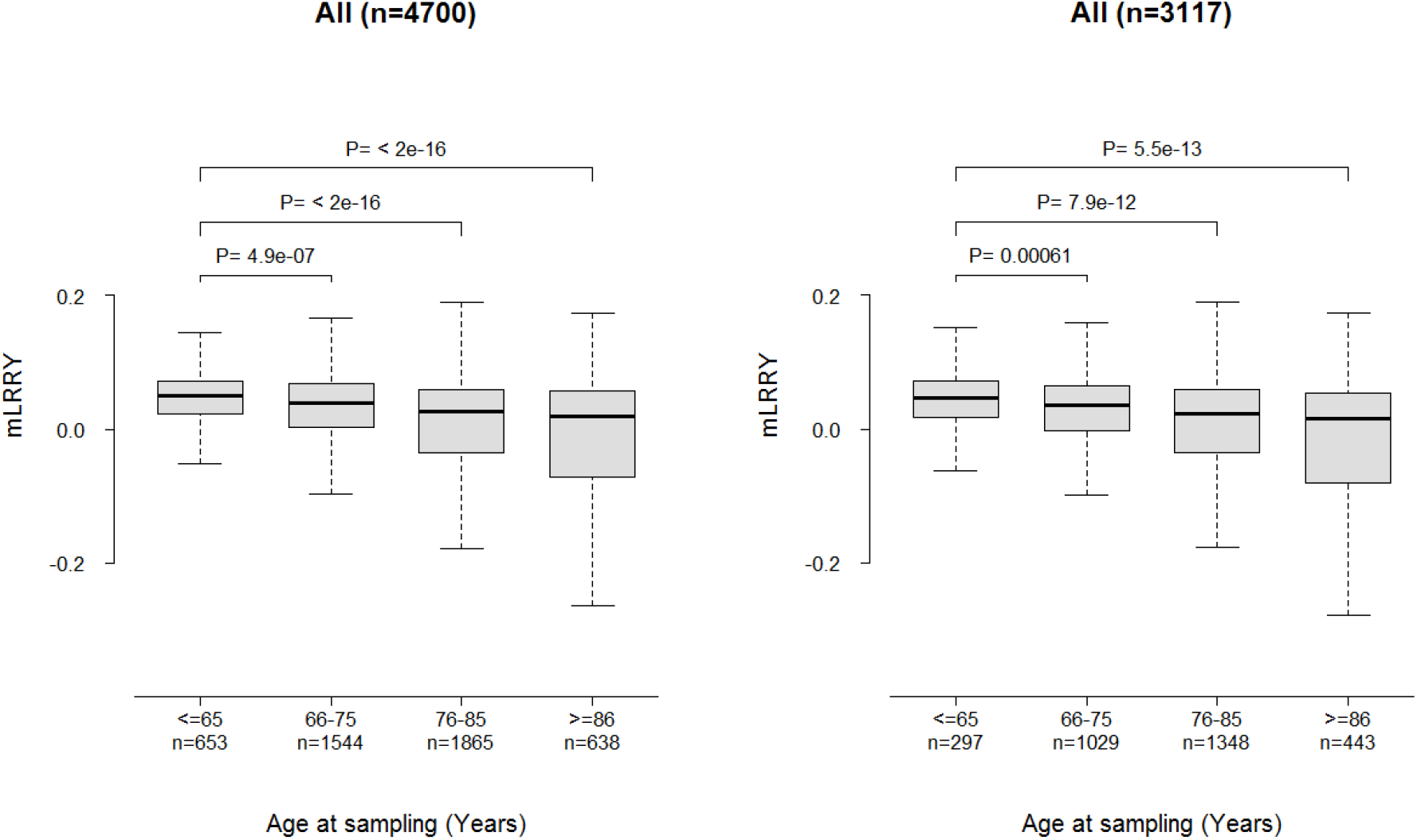
Normalized signal intensity of Y chromosome probe intensity by age at sampling. The median log R ratio difference between probes for chromosome Y and chromosome 1 (mLRRY) is plotted for four age ranges. Reported p-values are from Wilcoxan rank sum tests between indicated age groups. All individuals with available genotype array data are shown in the left figure, while individuals with genotype array data and an AD diagnosis are shown on the right.

We next performed Kolmogorov-Smirnov (KS) analyses examining differences in continuous mLOY between cases and controls. We further adjusted these analyses for age at sampling using Analysis of Covariance (ANCOVA). The KS test was significant (p = 0.0063) with AD cases having lower mLRRY than controls. ANCOVA analyses adjusting for age at sampling were significant only in the ADC3 cohort with no significant differences overall (p = 0.70).

### Association between LOY and Risk of AD Diagnosis in Follow-up

To evaluate the association between LOY and risk of developing AD in follow-up, we fit a Cox proportional hazards model associating binary LOY to years of disease-free follow-up within 1447 individuals for whom blood was sampled prior to any AD diagnosis. Of these, there were 42 subsequent AD events with all others remaining controls. Overall, we find a significantly increased hazard (1.98 [1.17–6.91], p=0.0021) in men with LOY versus those without (**Figure 2**).

**Figure 2.**
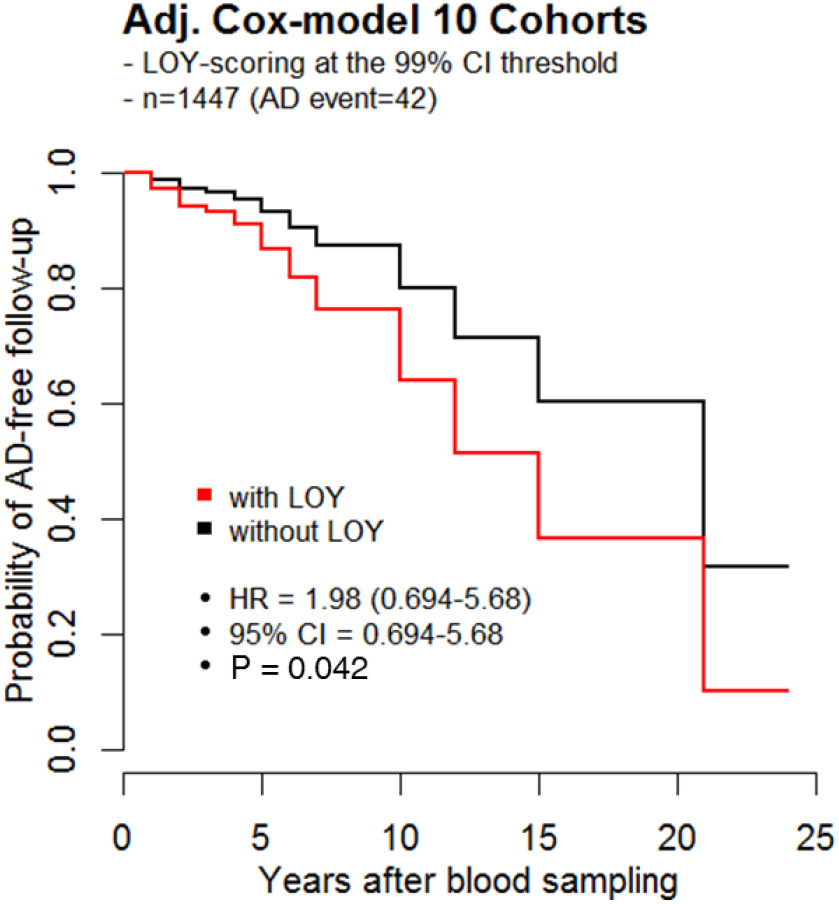
Cox Proportional Hazards Model of Incident AD Cases. Of 1447 individuals across 10 cohorts with blood samples drawn prior to AD diagnosis, a binary LOY threshold (using the 99% CI approach) shows a significant difference in probability of AD-free follow-up.

### Confirmation with Mendelian Randomization

To account for potential biases in sample ascertainment with respect to age and case/control status, we performed Mendelian Randomization analyses of LOY. To construct a genetic instrument, we considered significant SNPs from two published GWAS on LOY [12, 13]. Due to a large number of missing indels that were associated in Thompson et al., we restricted our analysis to an instrument using SNPs from Wright et al. using estimated betas and standard errors for LOY within our dataset. Previously published betas have a strong positive correlation with betas estimated in our LOY subset, with an R^2^ of 0.5167. As these results demonstrate the validity of our instrumental variable, we next compared these effects to estimated betas and standard errors for these same SNPs on AD case/control status in all males of the full ADGC dataset (n=10,013 across 26 cohorts, **Figure 3a**). Effect estimates for AD from these analyses were adjusted for age at onset/last follow-up, APOE status, and ancestry-based principal components.

**Figure 3.**
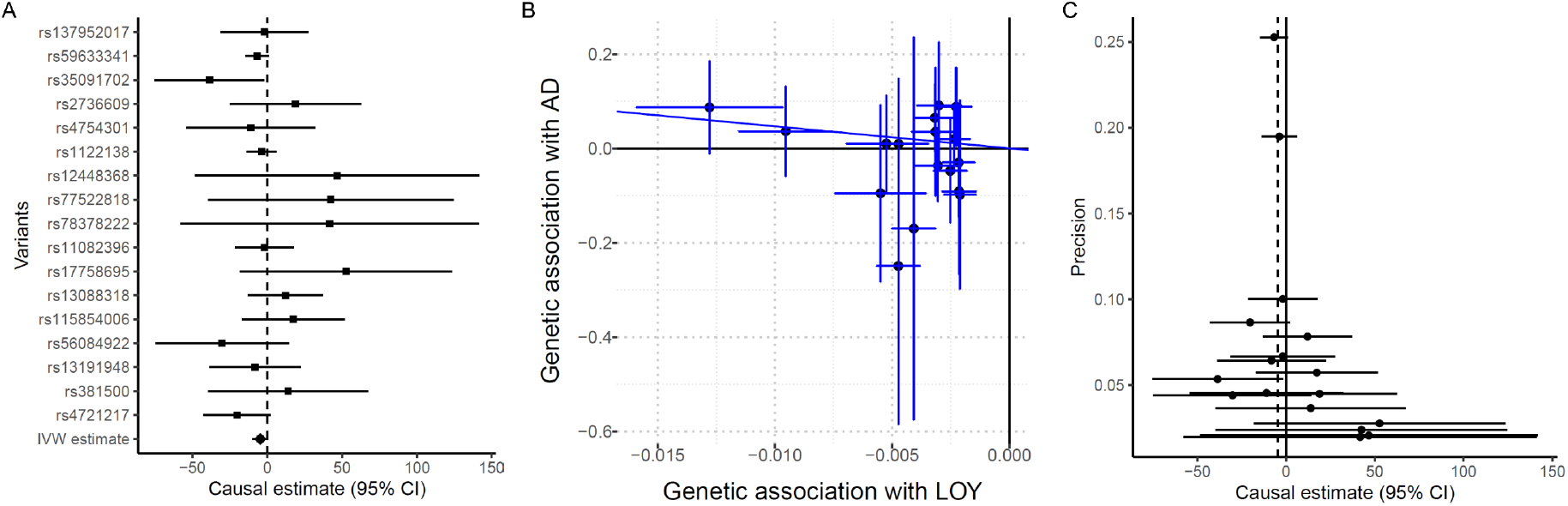
Two-sample Mendelian Randomization of LOY on AD. A) AD effect estimates for 10,013 males across 26 cohorts were generated for 17 SNPs from the Wright et al. instrument and compared to LOY effect estimates using the Inverse Variance Weighted (IVW) approach. B) Regression plot demonstrating correlation between effect estimates of AD and LOY assessed as Y chromosome probe intensity; decreasing Y intensity (or increasing risk of LOY) is associated to increased risk of AD. C) Funnel plot of variant-specific MR causal estimates against their precision, which is generally symmetrical, indicating that the overall casual estimates are less likely to be biased.

Using the Wright-based instrument, we had a significant two-sample Mendelian randomization effect using the Inverse Variance Weighted approach (p = 0.008, **Figure 3b**). This analysis also confirms the direction of association, with increasing genetic burden for LOY also increasing risk for AD. Visual inspection of the funnel plot (**Figure 3c**) did not reveal obvious signs of systematic bias, especially for more precise variants.

## Discussion

In this study, we have examined the relationship between somatic loss of the Y chromosome in males and AD risk across multiple US-based cohorts. We further validate previous, consistent reports that the degree of LOY within the peripheral blood increases with age. We also confirm a previous estimate that males with LOY are at approximately two-fold increased risk for AD [11]. To expand our assessments beyond the direct prospective or co-incident measurement of LOY, we also conducted Mendelian randomization analyses using a GWAS-based instruments for predicting LOY [12]. This analysis confirms a significant association between LOY and AD in males via two-sample MR. Together, these results provide additional evidence that somatic loss of Y is a potential prognostic indicator of AD risk, and that LOY and AD may share variants and/or causal mechanisms in their pathogenesis accounting for age.

The datasets examined here as part of the ADGC represent the largest sample size of males from the US with available genotyping data and clinically adjudicated AD phenotypes. When examining male individuals with available Y chromosome probes, the sample size of our hazard regression analyses is equivalent to that reported in Dumanski et al., with a similar median age at blood collection. One notable difference is in the number of incident cases, which are somewhat lower in our study. This is due to the cross-sectional design of the datasets used in our study; the majority of AD cases had their blood drawn at or after their initial AD diagnosis, and many of the incident AD cases that power our statistical analyses were initially enrolled as controls and later convert to cases.

Mendelian Randomization analysis further supports a relationship between genetic variants predicting LOY and AD risk. The limitations and assumptions of MR analyses have been previously noted [19]. Our significant IVW MR statistic may be somewhat sensitive to horizontal pleiotropy (where one or more of the LOY variants influence AD risk via a mechanism other than directly through LOY), however prior studies and our hazard regression analyses already provide evidence of a temporal relationship between the onset of LOY and the development of AD. Furthermore, even if the LOY instrument variants contribute to AD risk through a underlying pleiotropic mechanism, the utility of LOY as a possible biomarker for AD onset remains.

A more notable weakness of our MR analyses is potential confounding by other unmeasured conditions. LOY associates with competing causes of mortality such as cancer risk [10] and cardiovascular disease [14, 20] and we are unable to adjust for these conditions within the examined cohorts due to limited data collection, however previous studies suggest that accounting for these competing risks would increase LOY effects rather than cause spurious associations [11]. As increasing age is a risk factor for both LOY and AD, a prospective cohort design is ideal to demonstrate that LOY precedes AD onset. Given the noted difficulties in prospective recruitment for AD [21] and the time horizon over which LOY shows its effect, additional epidemiological confirmation of this effect may prove elusive. These results together with prior epidemiological and molecular studies confirm that somatic loss of Y is a potentially important prognostic marker of AD risk.

## Acknowledgements

The National Institutes of Health, National Institute on Aging (NIH-NIA) supported this work through the following grants: ADGC, U01 AG032984, RC2 AG036528; Samples from the National Cell Repository for Alzheimer’s Disease (NCRAD), which receives government support under a cooperative agreement grant (U24 AG21886) awarded by the National Institute on Aging (NIA), were used in this study.

We thank contributors who collected samples used in this study, as well as patients and their families, whose help and participation made this work possible; Data for this study were prepared, archived, and distributed by the National Institute on Aging Alzheimer’s Disease Data Storage Site (NIAGADS) at the University of Pennsylvania (U24-AG041689-01); NACC, U01 AG016976; NIA LOAD, U24 AG026395, R01AG041797; Banner Sun Health Research Institute P30 AG019610; Boston University, P30 AG013846, U01 AG10483, R01 CA129769, R01 MH080295, R01 AG017173, R01 AG025259, R01AG33193; Columbia University, P50 AG008702, R37 AG015473; Duke University, P30 AG028377, AG05128; Emory University, AG025688; Group Health Research Institute, UO1 AG006781, UO1 HG004610, UO1 HG006375; Indiana University, P30 AG10133; Johns Hopkins University, P50 AG005146, R01 AG020688; Massachusetts General Hospital, P50 AG005134; Mayo Clinic, P50 AG016574; Mount Sinai School of Medicine, P50 AG005138, P01 AG002219; New York University, P30 AG08051, UL1 RR029893, 5R01AG012101, 5R01AG022374, 5R01AG013616, 1RC2AG036502, 1R01AG035137; Northwestern University, P30 AG013854; Oregon Health & Science University, P30 AG008017, R01 AG026916; Rush University, P30 AG010161, R01 AG019085, R01 AG15819, R01 AG17917, R01 AG30146; TGen, R01 NS059873; University of Alabama at Birmingham, P50 AG016582; University of Arizona, R01 AG031581; University of California, Davis, P30 AG010129; University of California, Irvine, P50 AG016573; University of California, Los Angeles, P50 AG016570; University of California, San Diego, P50 AG005131; University of California, San Francisco, P50 AG023501, P01 AG019724; University of Kentucky, P30 AG028383, AG05144; University of Michigan, P50 AG008671; University of Pennsylvania, P30 AG010124; University of Pittsburgh, P50 AG005133, AG030653, AG041718, AG07562, AG02365; University of Southern California, P50 AG005142; University of Texas Southwestern, P30 AG012300; University of Miami, R01 AG027944, AG010491, AG027944, AG021547, AG019757; University of Washington, P50 AG005136; University of Wisconsin, P50 AG033514; Vanderbilt University, R01 AG019085; and Washington University, P50 AG005681, P01 AG03991.

The Kathleen Price Bryan Brain Bank at Duke University Medical Center is funded by NINDS grant # NS39764, NIMH MH60451 and by Glaxo Smith Kline. Genotyping of the TGEN2 cohort was supported by Kronos Science. The TGen series was also funded by NIA grant AG041232 to AJM and MJH, The Banner Alzheimer’s Foundation, The Johnnie B. Byrd Sr.

Alzheimer’s Institute, the Medical Research Council, and the state of Arizona and also includes samples from the following sites: Newcastle Brain Tissue Resource (funding via the Medical Research Council, local NHS trusts and Newcastle University), MRC London Brain Bank for Neurodegenerative Diseases (funding via the Medical Research Council),South West Dementia Brain Bank (funding via numerous sources including the Higher Education Funding Council for England (HEFCE), Alzheimer’s Research Trust (ART), BRACE as well as North Bristol NHS Trust Research and Innovation Department and DeNDRoN), The Netherlands Brain Bank (funding via numerous sources including Stichting MS Research, Brain Net Europe, Hersenstichting Nederland Breinbrekend Werk, International Parkinson Fonds, Internationale Stiching Alzheimer Onderzoek), Institut de Neuropatologia, Servei Anatomia Patologica, Universitat de Barcelona.

ADNI data collection and sharing was funded by the National Institutes of Health Grant U01 AG024904 and Department of Defense award number W81XWH-12-2-0012. ADNI is funded by the National Institute on Aging, the National Institute of Biomedical Imaging and Bioengineering, and through generous contributions from the following: AbbVie, Alzheimer’s Association; Alzheimer’s Drug Discovery Foundation; Araclon Biotech; BioClinica, Inc.; Biogen; Bristol-Myers Squibb Company; CereSpir, Inc.; Eisai Inc.; Elan Pharmaceuticals, Inc.; Eli Lilly and Company; EuroImmun; F. Hoffmann-La Roche Ltd and its affiliated company Genentech, Inc.; Fujirebio; GE Healthcare; IXICO Ltd.; Janssen Alzheimer Immunotherapy Research & Development, LLC.; Johnson & Johnson Pharmaceutical Research & Development LLC.; Lumosity; Lundbeck; Merck & Co., Inc.; Meso Scale Diagnostics, LLC.; NeuroRx Research; Neurotrack Technologies; Novartis Pharmaceuticals Corporation; Pfizer Inc.; Piramal Imaging; Servier; Takeda Pharmaceutical Company; and Transition Therapeutics.

The Canadian Institutes of Health Research is providing funds to support ADNI clinical sites in Canada. Private sector contributions are facilitated by the Foundation for the National Institutes of Health (www.fnih.org).

The grantee organization is the Northern California Institute for Research and Education, and the study is coordinated by the Alzheimer’s Disease Cooperative Study at the University of California, San Diego. ADNI data are disseminated by the Laboratory for Neuro Imaging at the University of Southern California. We thank Drs. D. Stephen Snyder and Marilyn Miller from NIA who are ex-officio ADGC members.

Support was also from the Alzheimer’s Association (LAF, IIRG-08-89720; MP-V, IIRG-05-14147) and the US Department of Veterans Affairs Administration, Office of Research and Development, Biomedical Laboratory Research Program. P.S.G.-H. is supported by Wellcome Trust, Howard Hughes Medical Institute, and the Canadian Institute of Health Research.

