## Supplemental Material for "Somatic Loss of the Y Chromosome and Alzheimer’s Disease Risk"

1) Cleveland Institute for Computational Biology, Department of Population and Quantitative Health Sciences, Case Western Reserve University, Cleveland, OH. 2) Department of Biostatistics, Epidemiology and Informatics, University of Pennsylvania, Philadelphia, Pennsylvania, USA. 3) John P. Hussman Institute for Human Genomics, University of Miami Miller School of Medicine, Miami, Florida, USA. 4) John T. Macdonald Foundation Department of Human Genetics, University of Miami Miller School of Medicine, Miami, Florida, USA. 5) Department of Immunology, Genetics and Pathology, Science for Life Laboratory, Uppsala University, 75108 Uppsala, Sweden. 6) Department of Neurology, University of Miami Miller School of Medicine, Miami, FL, USA.

**Alzheimer's Disease Genetics Consortium (ADGC)**

The ADGC dataset comprises multiple cohorts. In this study, participants from ten studies were included based on the Y chromosome coverage of their original genotyping. Cohorts included were the Adult Changes in Thought (ACT)/ Electronic Medical Records and Genetics (eMERGE) Study, the National Institute on Aging (NIA) Alzheimer Disease Centers (ADC3 through 7), the Chicago Health and Aging Project (CHAP), the Rush University Religious Orders Study/Memory and Aging Project (ROSMAP), and the Washington Heights-Inwood Community Aging Project (WHICAP). All analyses were restricted to individuals of European ancestry because there was an insufficient number of subjects from other ancestry groups to obtain meaningful results. All subjects were recruited under protocols approved by the appropriate Institutional Review Boards.

The ACT/eMERGE Studies (ACT): The ACT cohort is an urban and suburban elderly population from a stable HMO that includes 2,581 cognitively intact subjects age ≥ 65 who were enrolled between 1994 and 199820,21. An additional 811 subjects were enrolled in 2000-2002 using the same methods except oversampling clinics with more minorities. More recently, a Continuous Enrollment strategy was initiated in which new subjects are contacted, screened and enrolled to keep 2000 active at-risk person-years accruing in each calendar year. This resulted in an enrollment of 4,146 participants as of May 2009. All clinical data are reviewed at a consensus conference. Dementia onset is assigned half way between the prior biennial and the exam that diagnosed dementia. Enrollment for eMERGE Study began in 2007. A waiver of consent was obtained from the IRB to enroll deceased ACT participants. In total, ACT/eMERGE contributed data on 566 individuals with probable or possible AD (70 with autopsy-confirmation) and on 1,696 CNEs (155 with autopsy-confirmation) who were included in the analyses.

The NIA ADC Samples (ADC): The NIA ADC3 cohort included subjects ascertained and evaluated by the clinical and neuropathology cores of the 29 NIA-funded ADCs. Data collection is coordinated by the National Alzheimer's Coordinating Center (NACC). NACC coordinates collection of phenotype data from the 29 ADCs, cleans all data, coordinates implementation of definitions of AD cases and controls, and coordinates collection of samples. The ADC cohort consists of 2,288 autopsy-confirmed and 913 clinically-confirmed AD cases, and 519 cognitively normal elders (CNEs) with complete neuropathology data who were older than 60 years at age of death, and 744 living CNEs evaluated using the Uniform dataset (UDS) protocol2,3 who were documented to not have mild cognitive impairment (MCI) and were between 60 and 100 years of age at assessment.

Based on the data collected by NACC, the ADGC Neuropathology Core Leaders Subcommittee derived inclusion and exclusion criteria for AD and control samples. All autopsied subjects were age ≥ 60 years at death. AD cases were demented according to DSM-IV criteria or Clinical Dementia Rating (CDR) ≥ 1. Based on the data collected by NACC, the ADGC Neuropathology Core Leaders Subcommittee derived inclusion and exclusion criteria for AD and control samples. All autopsied subjects were age ≥ 60 years at death. AD cases were demented according to DSM-IV criteria or Clinical Dementia Rating (CDR) ≥ 1. Neuropathologic stratification of cases followed NIA/Reagan criteria explicitly, or used a similar approach when NIA/Reagan criteria4 were coded as not done, missing, or unknown. Cases were intermediate or high likelihood by NIA/Reagan criteria with moderate to frequent amyloid plaques5 and neurofibrillary tangle (NFT) Braak stage of III-VI6,7. Persons with Down's syndrome, non-AD tauopathies and synucleinopathies were excluded. All autopsied controls had a clinical evaluation within two years of death. Controls did not meet DSM-IV criteria for dementia, did not have a diagnosis of mild cognitive impairment (MCI), and had a CDR of 0, if performed. Controls did not meet or were low-likelihood AD by NIA/Reagan criteria, had sparse or no amyloid plaques, and a Braak NFT stage of 0 – II.

ADCs sent frozen tissue from autopsied subjects and DNA samples from some autopsied subjects and from living subjects to the ADCs to the National Cell Repository for Alzheimer's Disease (NCRAD). DNA was prepared by NCRAD for genotyping and sent to the genotyping site at Children's Hospital of Philadelphia. ADC samples were genotyped and analyzed in separate batches. ADC1 and ADC2 contributed 2,304 AD cases (1,761 autopsy-confirmed; 543 clinically-confirmed) and 675 CNEs (515 autopsy-confirmed; 160 clinically-confirmed), of which 1,595 autopsied-confirmed AD cases and 132 CNEs were analyzed in our previous study1. The ADC3 dataset contains 897 clinically-identified living cases (527 with autopsy-confirmation) and 588 CNEs (4 with autopsy-confirmation) who were genotyped between July and August 2010.

Alzheimer Disease Center (ADC) waves 4, 5, 6 and 7 (ADC4, ADC5, ADC6, ADC7): The ADC cohorts included subjects ascertained and evaluated by the clinical and neuropathology cores of the 29 NIA-funded ADCs. Data collection is coordinated by the National Alzheimer's Coordinating Center (NACC). NACC coordinates collection of phenotype data from the 29 ADCs, cleans all data, coordinates implementation of definitions of AD cases and controls, and coordinates collection of samples. ADC autopsied subjects have detailed neuropathological data and were older than 60 years at the time of death. Diagnoses for 61 AD cases and 23 controls in these datasets were confirmed pathologically Living ADC subjects were evaluated using the Uniform dataset (UDS) protocol2,3 and were included in the study if they were documented to not have mild cognitive impairment (MCI) and were between 60 and 100 years of age at the time of the most recent assessment. AD cases met NINCDS-ADRDA criteria for definite or probable AD, or had a Clinical Dementia Rating (CDR) ≥ 1. The 803 cases and 1,220 controls genotyped as part of ADC waves 4, 5, and 6 were genotyped on the Illumina HumanOmniExpress-24 beadchip.

The CHAP Project (CHAP): Chicago Health and Aging Project (CHAP): CHAP is an on-going community based study of individuals from a geographically defined community of 3 neighborhoods in Chicago, Illinois (Morgan Park, Washington Heights, and Beverly), with 6,158 participants in the first phase of the study (78.7% overall; 80.5% of the blacks, 74.6% of the whites) [citation- PMID: 14646025]. Data were collected in cycles of approximately 3 years; each consisting of an in-home interview of all participants and clinical evaluation of a random, stratified sample. The baseline cycle measured disease prevalence and provided risk factor data prior to incident disease onset. A cohort of 3,838 persons free of AD was identified; 729 persons were sampled for baseline clinical evaluation. Persons in the disease-free cohort had either good cognitive function at baseline, or if cognitive function was intermediate or poor, were free from AD at the baseline clinical evaluation. This disease-free cohort was evaluated for incident disease after an average of 4.1 years. Sampling for incident clinical evaluation was based on age, sex, race, and change in cognitive function (i.e., stable or improved, small decline, or large decline). The sample set available in the ADGC for genetic analyses included 32 AD cases and 197 persons free of AD at time of last assessment (all subjects were age 65 years or older at last assessment).

The ROS/MAP Studies: ROS/MAP are two community-based cohort studies. The ROS has been on-going since 1993, with a rolling admission. Through July of 2010, 1,139 older nuns, priests, and brothers from across the United States initially free of dementia who agreed to annual clinical evaluation and brain donation at the time of death completed their baseline evaluation. The MAP has been on-going since 1997, also with a rolling admission. Through July of 2010, 1,356 older persons from across northeastern Illinois initially free of dementia who agreed to annual clinical evaluation and organ donation at the time of death completed their baseline evaluation. Details of the clinical and neuropathologic evaluations have been previously reported26-29. A total of 1,072 persons passed genotyping QC. Of these, 296 met clinical criteria for AD at the time of their last clinical evaluation or time of death and met neuropathologic criteria for AD for those on whom neuropathologic data were available, and 776 were without dementia or MCI at the time of their last clinical evaluation or time of death and did not meet neuropathologic criteria for AD for those on whom neuropathologic data were available.

Washington Heights-Inwood Community Aging Project (WHICAP): WHICAP is a community-based longitudinal study of aging and dementia among elderly, urban-dwelling residents26,27. Beginning enrolment in 1989, WHICAP has followed more than 5,900 residents over 65 years of age, including white, African American, and Hispanic participants. Detailed clinical assessments were performed at approximately 24-month intervals over the 7 years of the initial study. All interviews were conducted in either English or Spanish. The choice of language was decided by the subject in order to ensure the best performance, and the majority of assessments were performed in the subject’s home, which included medical, neurological, and neuropsychological evaluations. Results of the neurological, psychiatric and neuropsychological assessments were reviewed in a consensus conference comprised of neurologists, psychiatrists, and neuropsychologists. Based on this review all participants were assigned to one of three categories: dementia, cognitive impairment or normal cognitive function. The sample set available in the ADGC for genetic analyses included 73 AD cases and 570 subjects with normal cognitive function.
